# Automated quantification of ecological interactions from video

**DOI:** 10.1101/2025.08.05.668677

**Authors:** Mariana Sadde, Benjamin T. Martin

## Abstract

1. Ecological interactions, such as predation, are fundamental events that underlie the flow and distribution of energy through food webs. Yet, directly measuring interaction rates in nature and how they vary across space and time remains a core challenge in ecology.
2. To address this, we developed a machine learning pipeline that combines object detection, tracking, behavioural classification, and bias correction to automatically quantify interaction rates from video. We applied this pipeline to estimate feeding rates of the planktivorous reef fish *Chromis multilineata* in situ.
3. We show that the tool generates precise, unbiased estimates of planktivory at fine temporal and spatial scales, and use it to reveal how feeding rate changes in response to predator presence and proximity to refuge.
4. More broadly, this method provides a scalable, non-invasive framework for quantifying interactions in the wild, enabling new opportunities to test ecological theory and quantify energetic fluxes in nature.

## Introduction

The dynamics of ecological systems emerge from vast numbers of individual interactions among predators and prey, competitors, and mutualists. However, directly quantifying the *rates* of these interactions and how they vary across environmental conditions in natural systems remains a major challenge. This difficulty stems largely from the logistical and methodological constraints of observing interactions as they happen: they are often fleeting or cryptic, making them hard to capture using traditional field techniques [1].

As a result, ecologists have relied on a range of time-limited or indirect methods to estimate interaction rates. Focal observations [2], camera traps [3], and tethering or exclosure experiments [4] have all been used to document interactions, but are typically constrained in spatial or temporal scale. Stable isotope analysis [5], gut content examination [6], and DNA metabarcoding [7] can identify trophic links, but offer only coarse, time-averaged measurements that do not resolve the frequency or context of interactions. Species removal experiments [8] and co-occurrence-based network inference [9] similarly infer interactions from outcomes or associations, rather than direct measurements of interactions. These approaches have advanced our understanding of who interacts with whom, but have limited capacity to quantify *how often* interactions occur, how they are distributed in space and time, or how they respond to changing environmental conditions.

One promising avenue for directly quantifying the rate of ecological interactions in the field is the use of machine-learning-based computer vision tools. In particular, deep-learning-based computer vision allows for automated detection, classification, and tracking of objects in both images and video [10,11], opening the door to continuous, high-throughput observation of ecological processes. Although deep-learning has already been applied in species detection, individual identification, and trait estimation, its use for detecting *interactions* between organisms, especially quantifying their frequency over time, remains limited [12,13]. Developing tools that leverage computer vision to directly and continuously measure interaction rates offers a powerful opportunity to overcome long-standing challenges in ecological field studies.

In this study, we develop, test, and apply a machine-learning-based computer vision pipeline to quantify feeding rates of *Chromis multilineata* (brown chromis), a common, planktivorous damselfish on Caribbean reefs [14], and investigate how these rates are modulated by predator presence and proximity to refugia. Planktivorous fishes, such as brown chromis, play a critical role within coral reefs, capturing allochthonous, current-derived zooplankton and converting it into reef-based biomass and metabolic byproducts that contribute to nutrient cycling, and support coral, invertebrates, and other fish [15–18]. Although factors that influence planktivorous fish feeding rates, and, hence, pelagic subsidy influxes into coral reefs, are largely unknown [15–17], behavioural trade-offs planktivores experience in response to predation risk—known as non-consumptive effects (NCEs) [19–24]—may significantly alter the rate and spatial distribution of planktivory [15]. Such NCEs can cause prey to reduce foraging or increase refuge use in response to real or perceived threats [21,24–27], with ecosystem-scale consequences [19,20,25,28–32]. Understanding behavioural controls on planktivory may be especially important for degraded reefs, where external energy subsidies could help maintain fish biomass and ecosystem function in the absence of high coral cover [15]. More broadly, our approach demonstrates how machine-learning-based tools can help overcome long-standing challenges in field ecology by enabling direct, scalable, and behaviourally explicit measurements of species interactions, offering new insights into how ecological processes are structured across space, time, and environmental gradients.

## Methods

### Collection of brown chromis footage in the field

The data used for analysis comprised underwater videos of site-attached aggregations of brown chromis from multiple coral-head sites in a shallow reef flat off the coast of Cas Abao beach, Curacao (12.2331°N, –69.0972°W, at 3-4m depth), taken with GoPro 9 cameras (4K, 60fps, wide-angle setting). These videos were recorded from a custom-made underwater frame with a two-by-two-meter base placed directly on the sea floor around a coral-head site. Each frame was equipped with eight cameras, aimed inwards towards the coral head from the outer corners, with two cameras each, recording 50cm from the seafloor. For our analysis, we only used data from one camera per site, the one with the most comprehensive view of the chromis aggregation. Between November 15^th^ and December 6^th^, 2023, 12 diurnal deployments were made for approximately one hour each, two to four times per site. In addition, we used videos from short-term (5-10 minute) supplemental deployments at various coral colonies to generate additional images of chromis to train the object detection model.

### Overview of machine learning pipeline to estimate planktivory rate

Our machine learning pipeline was designed to detect and count the number of predatory strikes made by brown chromis aggregations per unit of time, to estimate per capita feeding rates of brown chromis (strikes s^-1^fish^-1^). To do this, we took advantage of the fact that brown chromis, like many other planktivorous damselfishes, rapidly extend their jaws to capture evasive zooplankton, primarily copepods [33]. Importantly, such jaw extensions are visible from videos of chromis taken at close range (1-2 meters) with a high-resolution camera.

The pipeline works in several steps. First, raw videos of brown chromis aggregations are fed into an object-detection and tracking algorithm to identify individual brown chromis in a video and track their position over time. This tracking information is then used to generate cropped videos of each detected and tracked chromis in a video (see **Supplemental Video 1**). Each of these cropped videos of individual chromis is then fed to a frame-by-frame behavioural classification model trained to detect predatory strikes. The average per-capita feeding rate of a brown chromis aggregation over a particular time period can then be calculated by dividing the number of feeding strikes at zooplankton by the amount of time chromis were oriented such that their strikes could be detected. In addition, we evaluated our model for biases in strike detection due to the apparent size, position or background colours of tracked chromis, and then corrected for these to generate unbiased strike rate estimates.

### Model training – Object detection

To train a model to recognise brown chromis, bounding box annotations were made of individuals within still images of videos collected in the field during the deployment period, using CVAT [34]. A “YOLOv8 large” model was trained on the annotated dataset, comprising 1 036 images, for 400 epochs [35].

### Model training – Image classification

The trained object detection model described above was used with bot-sort, a tracking algorithm [36], to track the movement of individual brown chromis over time. This resulted in a dataset with the locations of each tracked chromis across the frames in which they were tracked. We used this dataset to extract tracked chromis videos, cropped into 220×220 pixels and centred on the midpoint of their respective bounding box pixel coordinates (see **Figure 1, Supplemental Video 1**). In cases where the bounding boxes of tracked chromis were larger than the cropped region dimensions, the bounding box image was downsized so that the dimensions of all tracked chromis videos were identical.

**Figure 1.**
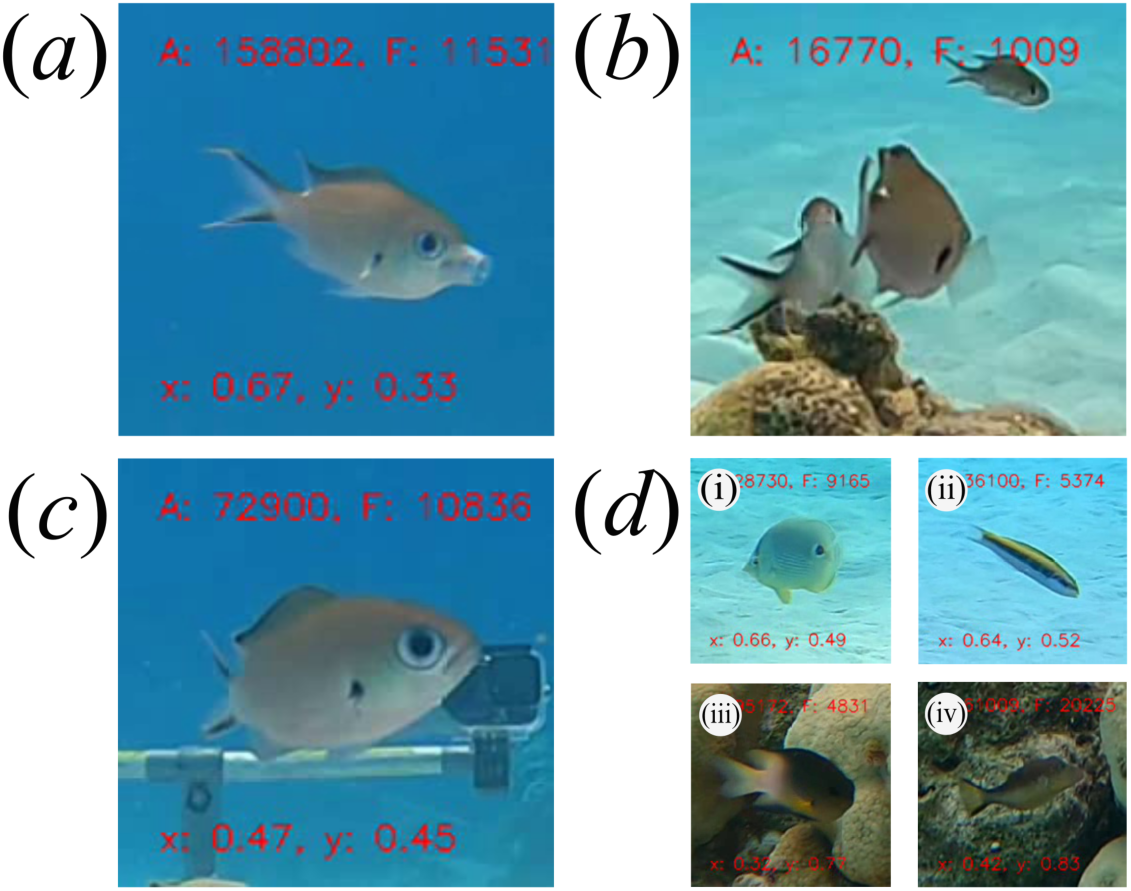
Individual brown chromis behaviour classes used for training the image classification model. (a) Feeding, (b) mouth not visible, (c) not feeding mouth visible, (d) (i-iv) non-chromis.

We wanted to calculate mean per-capita feeding rates as a function of time. However, whether or not a brown chromis is striking at prey can only be determined in periods when the chromis is oriented such that its mouth is visible. Additionally, in some instances, due to errors in the object detection model, non-chromis fish were tracked, which are not of interest to our study. Therefore, each frame of the tracked chromis video clips was annotated as one of four classes: “feeding”, “not feeding mouth visible”, “mouth not visible” and “non-chromis” (see **Figure 1**). We labelled each frame of the tracked fish videos using a custom annotation tool created in MATLAB [37]. To generate a training database for the frame-by-frame behavioural classification model, we extracted all “feeding” frames from the annotated video clips, and one of every five frames for each of the other classes. This resulted in a dataset comprising 173 813 images, divided within each class into training, validation and testing datasets under a 70%, 20%, 10% split, which was used to train the image classification model (“YOLOv8 large” classification model) for 100 epochs [35].

### Generating feeding-rate estimates for brown chromis aggregations

To generate per-capita feeding rate estimates for a brown chromis aggregation, we applied the detection, tracking, and behavioural classification steps described above. These model inference steps, as well as model training, were implemented in Python using Google Colab notebooks, executed on an NVIDIA A100 GPU. The model output at this stage provides predictions of the most likely behavioural class for each frame of each tracked chromis. To convert this to estimates of per-capita feeding rates, several post-processing steps were applied. First, tracked chromis clips shorter than or equal to 20 frames were excluded, as videos at or below this frame length are often part of the “non chromis” class, and we also excluded any track that had 10% or more of its frames with “non-chromis” labelled as the most likely class. This threshold was determined to exclude nearly all non-chromis tracks, while excluding relatively few true chromis tracks.

Secondly, because feeding strikes by chromis species typically last ∼30-60 milliseconds [33], we often observed 2-4 consecutive frames where the most likely class was “feeding”. We therefore counted such consecutive detections of feeding behaviours as one feeding event. Additionally, in rare cases, we observed prolonged jaw extensions, which appeared to be a form of stretching behaviour unrelated to feeding. We therefore treated periods of prolonged jaw extensions (> 5 frames) as non-feeding events, and relabelled these frames from “feeding” to “not feeding mouth visible”. After these post-processing steps, we calculate the per-capita feeding rate of chromis as the number of detected feeding events per amount of time when the mouths of chromis were labelled as visible i.e., sum of feeding frames and not feeding mouth visible frames.

### Model evaluation and bias correction

We evaluated the precision and recall of the **brown chromis detection model** on the “test” object detection dataset, comprising 149 images. We also evaluated the precision and recall of the **behaviour classification model** as described by Belcher *et al.* [11] on the classification test dataset comprising 24 837 images (557 “feeding”, 5 507 “mouth not visible”, 584 “non-chromis”, and 18 189 “not feeding mouth visible”).

In addition, we generated a dataset of 1 998 manually labelled videos of individually tracked brown chromis, randomly selected from all 12 deployments and not used for model training or validation. This dataset was used to assess bias in feeding rate estimates and evaluate the accuracy of feeding rate estimates. Classification errors (precision and recall < 1) can bias estimates; for example, recall of 0.5 for feeding events with perfect precision would lead to feeding rate estimates 50% lower than the true rate. Moreover, if model performance varies systematically across image features (e.g. bounding box size, background colour), it may bias comparisons of feeding rates across contexts. We therefore evaluated how recall and precision for the two behavioural classes relevant to feeding rate—“feeding” and “not feeding mouth visible”—varied with frame-level characteristics affecting image quality or visibility. These included bounding box area (larger for fish closer to the camera), x–y coordinates (higher image quality near the centre of the frame), and the median red, green, and blue (RGB) pixel values of the image background (see **Supplementary Figure 2**). Median RGB values for each frame were calculated from the outer row of pixels from the cropped images to capture background colour while excluding the fish.

To assess these sources of bias, we generated classification predictions for each frame in the 1 998 labelled videos and examined how precision and recall varied with the image-level covariates above. Full results are presented in **Supplementary Table 1**. The most important factor affecting recall was bounding box area: recall approached zero for small boxes and one for large bounding boxes. To avoid high error rates in feeding estimates, we excluded data with bounding box areas below the threshold where feeding recall dropped below 0.5 (<1 451 pixels). Thus, fish that are too far away from the camera, appearing too small in the image to reliably detect feeding events, are excluded from feeding rate calculations.

We then fit logistic regression models to predict precision and recall for the “feeding” and “not feeding mouth visible” classes as a function of frame-level variables, using the dredge function in the *MuMIn* package in R [38] and AIC to select the best models (**Supplementary Table 1**). These predictions were used to correct per-capita feeding rate estimates according to:

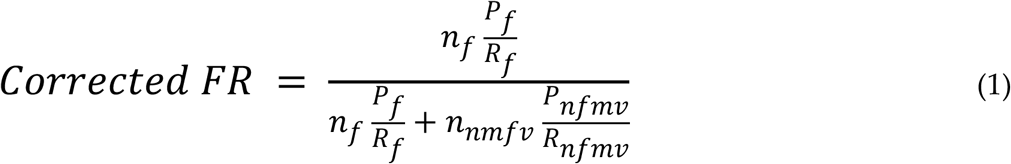

where *n_f_* and *n_nfmv_* are the number of predicted “feeding” and “not feeding mouth visible” frames, respectively, and *P* and *R* represent the average precision and recall for each class (indicated by subscripts) over the period in which feeding rate is calculated.

Finally, we evaluated the accuracy of corrected feeding rates as a function of the amount of data used in their calculation. We randomly sampled 10 000 subsets of the 1 998-video test set, ranging from 1 to 200 tracks. For each sample, we compared the corrected estimate to the ground-truth feeding rate and calculated the standard deviation of these differences across samples, binned by total duration (in 10-second intervals). This analysis allowed us to quantify how the precision of feeding rate estimates improves with increasing data.

### Applying the machine-learning tool

To demonstrate the utility of the pipeline, we used it to investigate predators’ influence on the feeding behaviour of brown chromis, in reactive and proactive predator avoidance contexts [25]. In the case of **reactive predator avoidance behaviours**, triggered by the presence of predators, we identified periods from the footage collected in the 12 deployments in which a predator, *Caranx ruber* (the bar jack), swam through the video-recording quadrat set-up, and observed how chromis groups reacted to this. The collective response of chromis to predator encounters was fairly stereotyped, allowing us to categorise chromis group behaviour into five states (“feeding normally” before and after predator disturbance, and well as “moving towards”, “hiding in”, and “moving away from” a coral refuge, see **Figure 2**). For each predator approach, we quantified the mean per-capita feeding rate in each of the five states. A Kruskal-Wallis test was conducted to test for significant differences in feeding rates between the five brown chromis group behaviours, followed by Dunn’s post-hoc test if significant. Individual track data from group behaviour phases in each predator encounter video were also grouped into disturbed (“moving towards”, “hiding in” and “moving away from” refuge) and undisturbed categories (“feeding normally” before and after predator disturbance), to calculate feeding rate for each, and test for significant differences between them using a paired t-test. We hypothesised that brown chromis would feed at reduced rates while disturbed by predators compared to undisturbed periods. Feeding rate estimates per behavioural phase for each video, calculated using equation (1), were treated as raw data points.

**Figure 2.**
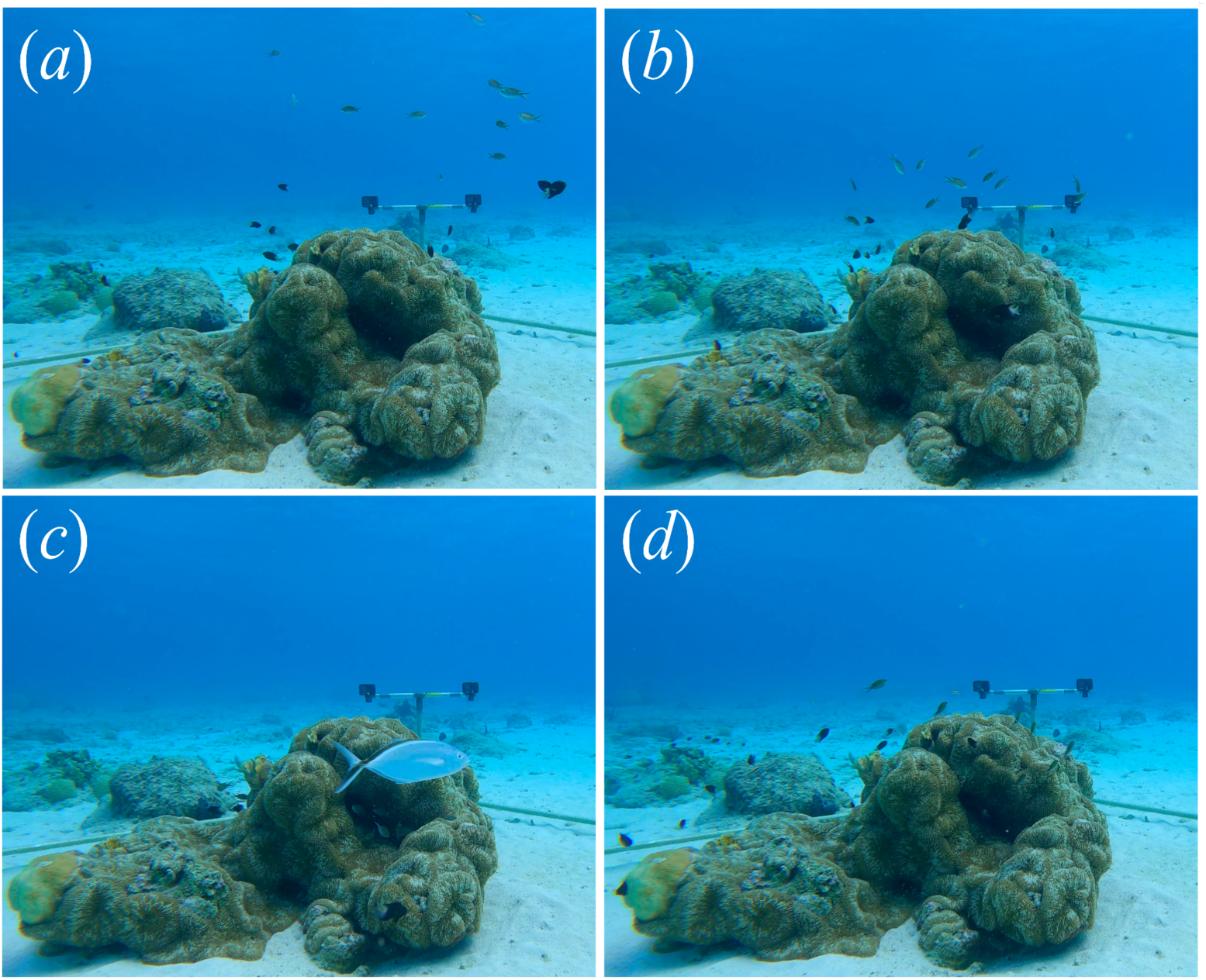
Brown chromis group behaviour categories. (a) Feeding normally (before or after a predator disturbance); (b) moving towards refuge; (c) hiding in refuge (with bar jack predator in view); (d) moving away from refuge.

**Figure 3.**
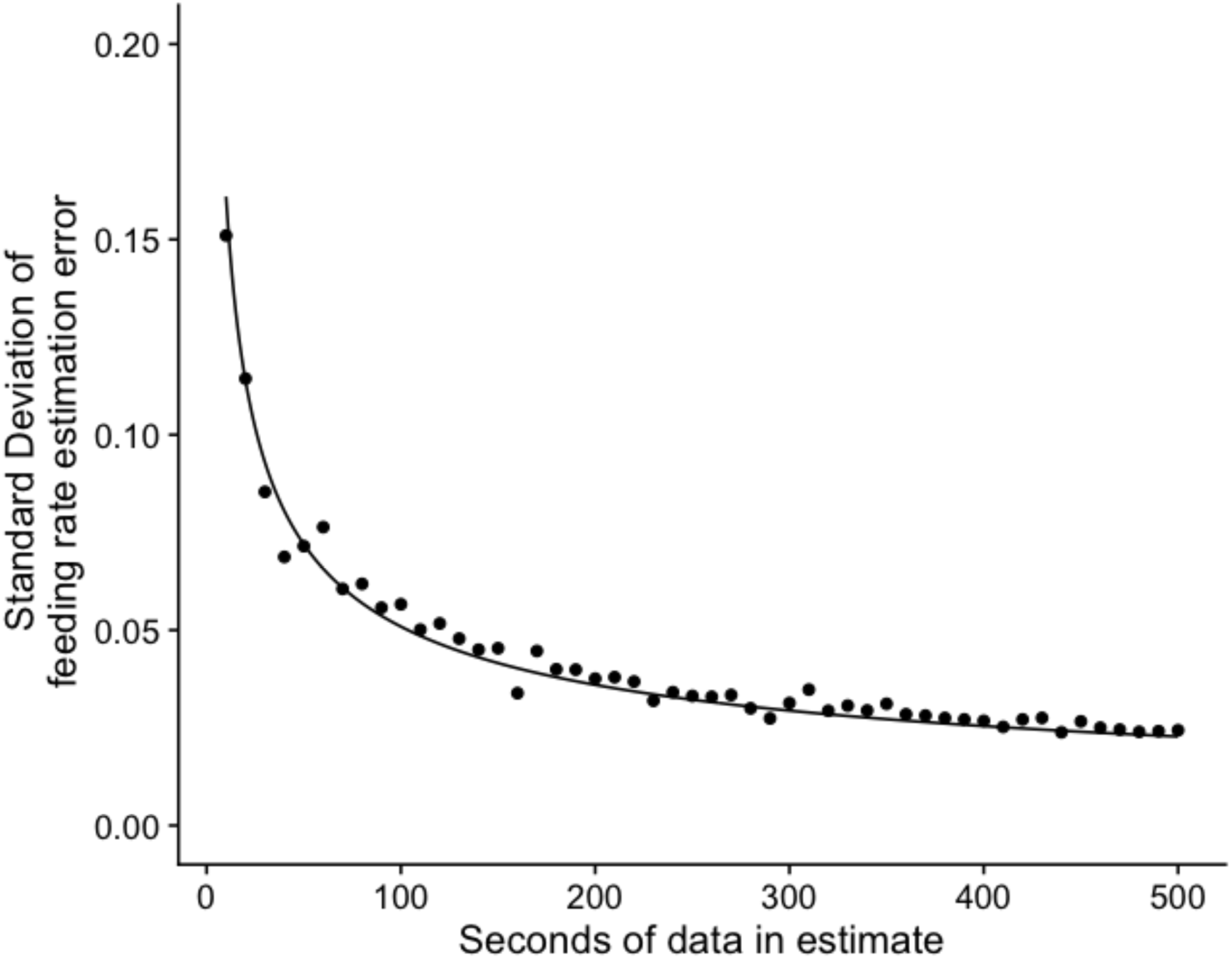
Precision of feeding rate estimates as a function of dataset size. Points depict the standard deviation of the differences between predicted and ground truth feeding rate estimates (SDE_FR_) as a function of dataset size (number of seconds of tracked fish in the estimate), binned into 10-second intervals (see “Model Evaluation and Bias Correction”). SDE_FR_ scaled with the inverse square root of dataset size (fitted line). For reference, the average per-capita feeding rate of brown chromis in our study was 0.54 (strikes s^-1^ fish^-1^).

In the case of proactive predator avoidance (i.e., behavioural responses related to proximity to refuge), we applied the machine-learning pipeline to all 12 one-hour deployments. We evaluated how feeding rates varied along the vertical axis of the water column, hypothesizing that chromis feed more when positioned farther from the coral refuge (i.e., higher in the water column). To test this, we grouped chromis detections within each deployment into quartiles based on their *y*-pixel (vertical) coordinates and estimated feeding rates for each quartile. To assess systematic variation in feeding activity across depths, we scaled the feeding rate in each quartile by dividing it by the mean feeding rate across all quartiles for that deployment. Thus, values greater than 1 indicate above-average feeding activity at a given depth, while values less than 1 indicate below-average feeding. These deployment-scaled feeding rates for each depth quartile were treated as raw data points. A one-way ANOVA was used to test for differences in deployment-scaled feeding rates between the four vertical space quantiles, followed by a Tukey post hoc test if significant.

All data analysis was conducted in R version 4.3.1 [39], using readxl, tidyverse, readr, car, cowplot, png, dunn.test, ggplot2, dplyr, tidyr, and patchwork packages [40–50].

## Results

### Model performance

Bounding box precision and recall values for the object detection model were 0.955 and 0.943, respectively, indicating most chromis in a video are detected and tracked. Errors in the chromis detection model generally only impact estimates of feeding rate by reducing the amount of data used for feeding rate calculations, as undetected and non-tracked chromis are excluded from these calculations.

Performance of the behavioural classification model was high overall. On the 1 998-track test dataset, the precision and recall for feeding events was 0.858 and 0.605, respectively. However, after excluding data with chromis bounding box sizes lower than 1 451 pixels, these values increased to 0.865 and 0.761. Consequently, most feeding events are successfully detected by the model with relatively few false positives. A full description of model performance for all behavioural classes is given in **Supplementary Tables 2-4**.

After applying the bias correction functions for precision and recall of the “feeding” and “not feeding mouth visible” eq. 1, we found that the precision of the feeding rate estimates scaled with the inverse square root of the amount of seconds of data used in the feeding rate calculation, such that standard deviation of feeding rate estimation error (SDE_FR_) was:

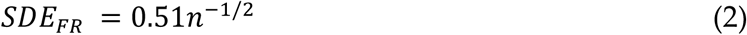

where n is the number of seconds of data ((“feeding” + “not feeding mouth visible frames”)/ 60) used in the feeding rate estimate. Thus, our model allows for accurate estimates of chromis strike rates at relatively high temporal resolution (10s to 100s seconds).

### Feeding behaviour of brown chromis

A total of 31 predator encounters were observed in the field-collected footage, 2.58 ± 2.02 (SD) times per deployment, and were used for analysis. The average total time of disturbance per predator encounter, or the total time spent “moving towards refuge”, “hiding in refuge” and “moving away from refuge”, was 22.71 ± 15.93 (SD) seconds. An example of model predictions of individual brown chromis fish detections and their behaviours, as well as the feeding rate through time across the different collective behaviours for a single predator encounter video, is shown in **Figure 4**.

**Figure 4.**
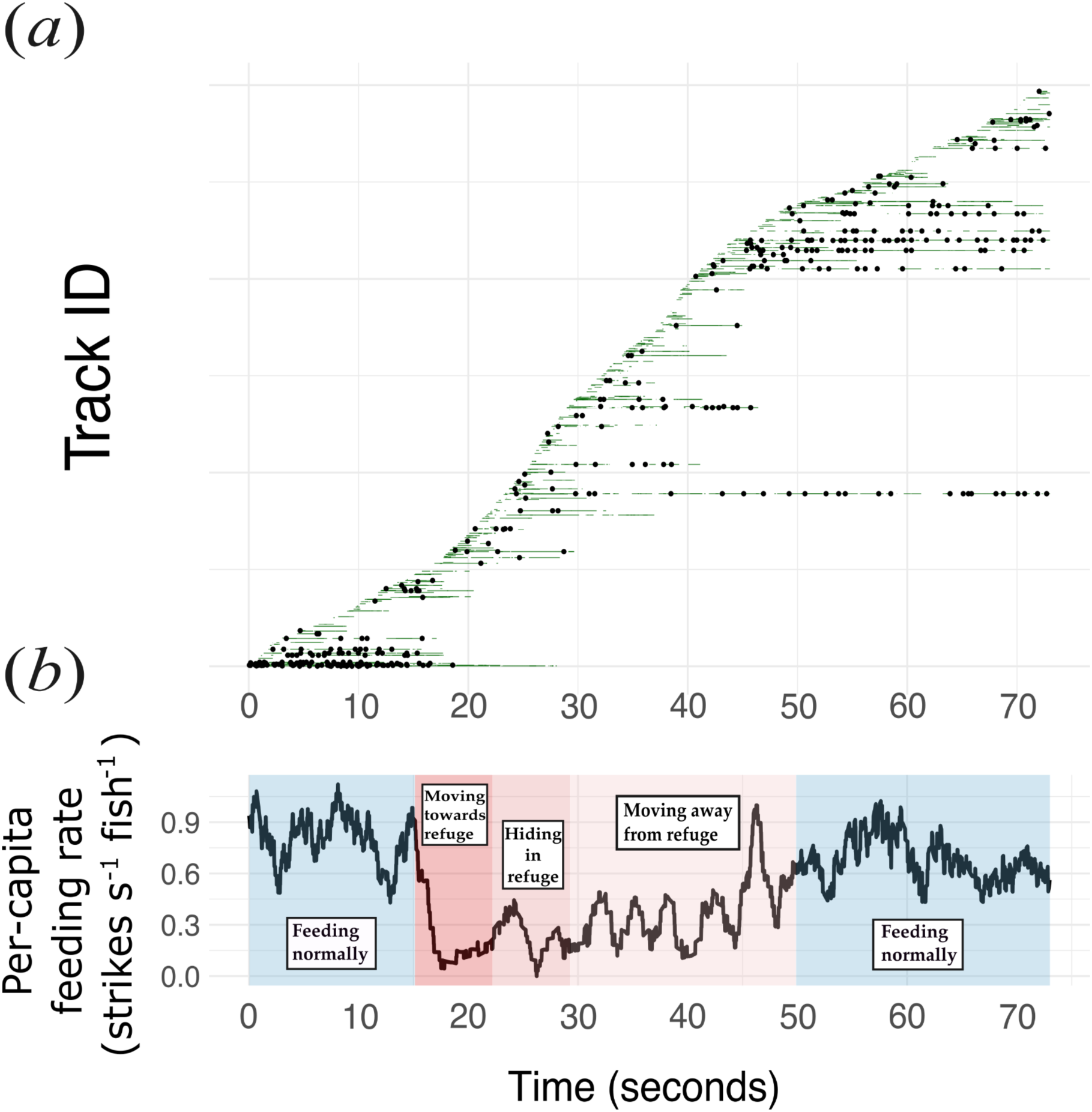
Feeding rates and behaviour of a brown chromis group and its individually tracked members within a predator encounter video. (a) Individual brown chromis tracks within a predator encounter video and behaviours predicted for these: horizontal lines represent individual tracks; white lines, green lines and black dots correspond to “mouth not visible”, “not feeding mouth visible” and “feeding” classes, respectively. (b) Average feeding rate per second (using an 8-second moving window) of a chromis aggregation through time across the five group behaviour phases.

#### Feeding rate during predator approaches

Feeding rates of chromis showed a systematic decrease in the presence of predators. Brown chromis aggregations had the highest feeding rates when “feeding normally”, before (mean (SE) = 0.56 (0.04) strikes s^-1^ fish^-1^), and after a predator disturbance (0.55 (0.04)), lower feeding rates when moving towards (0.33 (0.03)) or away from the refuge (0.36 (0.04)), and the lowest feeding rates when hiding in the refuge (0.21 (0.04)), though the latter were not significantly different from those exhibited when moving towards the refuge (Kruskal-Wallis: chi-squared = 46.518, df = 4, p < 0.0001), (Dunn’s post-hoc: p<0.05) (**Figure 5a**). Overall, the average feeding rate of brown chromis aggregations in the presence of predators (disturbed stages) was approximately half of that exhibited during undisturbed stages (t = 6.11, df = 29, p-value < 0.0001) (**Figure 5b**).

**Figure 5.**
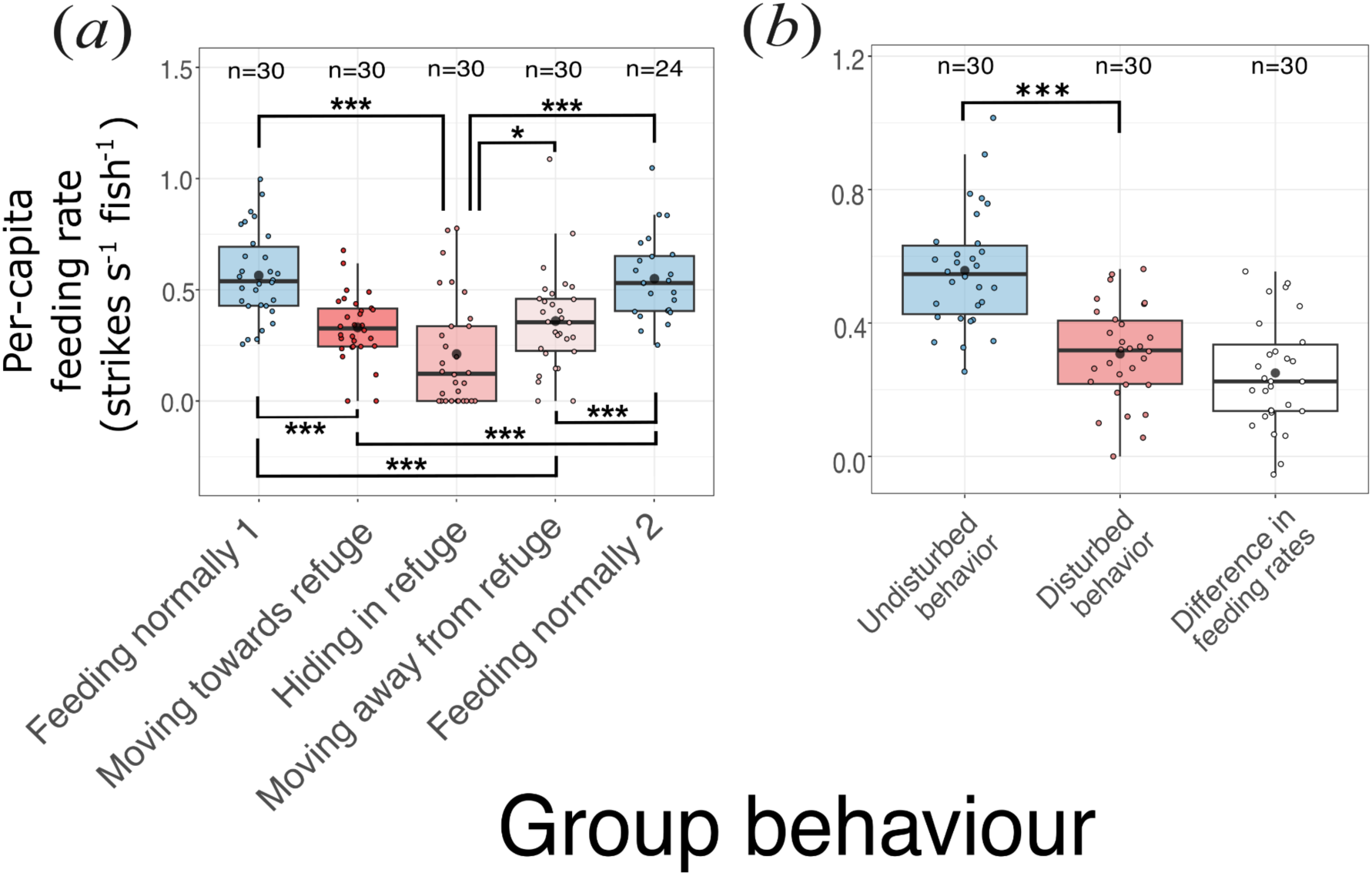
Feeding rate of brown chromis in response to predators. (a) Feeding rate of brown chromis aggregations per group behaviour phase during predator approaches. (b) Feeding rate during disturbed vs. undisturbed behaviours. Boxes (coloured by group behaviour) represent the interquartile range, horizontal black lines within correspond to medians, black dots to means, coloured dots to raw data points (coloured by group behaviour), sample size per group is displayed after “n=” above each box. Raw data points correspond to (within a predator encounter video): (a) average feeding rate for each behavioural phase; (b) average feeding rate for disturbed and undisturbed behaviours, and the difference between these. Whiskers extend to the maximum and minimum values within 1.5*IQR. Significant differences are indicated by “*” and “***” at p=0.01-0.05 and p=0.000-0.001 level, respectively.

#### Feeding rate of brown chromis across a vertical profile

Deployment-scaled feeding rate for brown chromis was significantly higher in upper levels of the water column (ANOVA: F=32.6; df=3, 44; p<0.0001), (**Figure 6**). Specifically, deployment-scaled feeding rate of brown chromis in the three upper quartiles was ∼20-30% higher than that of the lowest (Tukey HSD post-hoc: p<0.0001), and ∼10% higher for the second highest quartile compared to the third (Tukey HSD post-hoc: p=0.0472).

**Figure 6.**
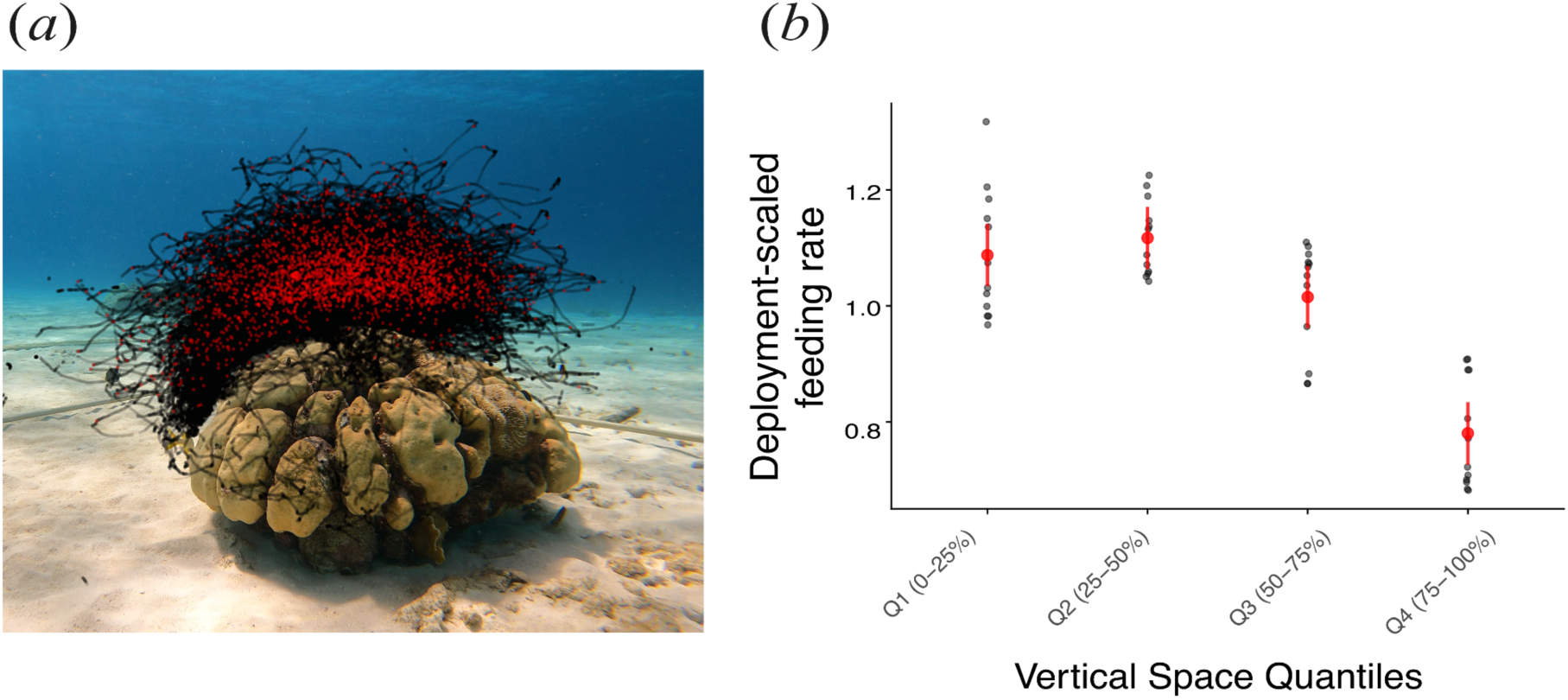
Feeding behaviour of individual brown chromis fish across a vertical profile. (a) Still image of coral-head site 2, with superimposed red and black points corresponding to x and y-axis coordinates of feeding and non-feeding events, respectively, of individual brown chromis within a one-hour deployment. (b) Deployment-scaled feeding rates (feeding rate of each quartile divided by the mean feeding rate across all depth quartiles for the deployment) across the depth quartiles (grey points), where Q1, Q2, Q3 and Q4 correspond to each quartile, ordered from highest to lowest position in the water column; red points and error bars correspond to predicted deployment-scaled means and 95% confidence intervals.

## Discussion

Ultimately, understanding how energy flows through ecological systems and predicting how such systems will respond to environmental change depends on knowing the rates at which species interact. While ecological theory often hinges on interaction rates (e.g. feeding, competition, mutualism), these rates are rarely measured directly in the field, especially at fine temporal and spatial scales. In this study, we developed and applied a machine learning-based computer vision pipeline that automates the detection of feeding behaviour in a coral reef planktivore, *Chromis multilineata*. Using over 12 hours of field video and more than 30 predator-prey encounters, we demonstrated that this tool can generate high-resolution, precise estimates of feeding rates, providing a scalable approach to quantify ecological interactions in situ.

### Model Performance

Importantly, the utility of tools for measuring interaction rates depends on their accuracy. We found that by explicitly modelling and correcting for sources of error in our behavioural classification model, we were able to produce precise and unbiased estimates of feeding rates at fine spatial and temporal scales. Although the classifier was not perfect (recall = 0.76 for feeding events), accounting for how recall and precision varied with frame-level characteristics allowed us to correct for systematic biases in feeding rate estimates. Because individual errors in detecting feeding events tend to average out over time, the accuracy of feeding rate estimates improves with the amount of data used. For example, just 60 seconds of cumulative tracked data (e.g., 10 fish tracked for 6 seconds each) yielded feeding rate estimates with a standard deviation of errors equivalent to 12% of the mean chromis feeding rate. With 600 seconds of tracked data, this error drops to just 4% of the mean. Thus, even relatively short observation periods can yield precise feeding estimates, and longer periods offer near-ground-truth accuracy, allowing for precise quantification of changes in the rates of ecological interactions over fine temporal and spatial scales.

### Findings from this application

We demonstrated the types of ecological questions that can be addressed with high-resolution measurements of interaction rates by analysing variation in brown chromis feeding behaviour in response to predation risk, both reactive, during predator encounters, and proactive, across vertical gradients of perceived threat. During predator encounters, feeding rates dropped by approximately 50%, consistent with our hypothesis that reactive predator avoidance behaviour reduces foraging opportunities. Specifically, chromis feeding rates were significantly higher during undisturbed periods, when fish were dispersed and feeding high in the water column, compared to periods when they were moving toward, hiding in, or moving away from the coral-head refuge. Notably, the reduction in feeding rates in the presence of predators (∼50%) was higher than the observed reduction in feeding for chromis distributed in the lowest depth quartile during the full deployments, suggesting the reduction in feeding in periods when a predator is present go beyond the expected reduction in feeding rates due to their proximity to the refuge alone. Perhaps more surprising is that chromis continue to feed at all while actively moving back to the refuge and hiding in close proximity to a predator. This suggests strong selective pressure on planktivores to maintain energy intake even in the face of elevated predation risk.

In addition to reactive responses to predator presence, we observed evidence of proactive risk avoidance, a spatial pattern in feeding behaviour that persisted even in the absence of nearby predators. Across all deployments, chromis consistently fed at higher rates when occupying upper regions of the water column. When in the lowest quartile of their distribution (i.e., closest to refuge), chromis fed at rates 20–30% lower than in higher strata. This reduction likely reflects both reduced access to zooplankton, which concentrate higher in the water column due to currents [16,51], and increased competition among more tightly aggregated individuals [27,52]. Given that predator approaches were relatively rare and short (∼twice per hour, lasting ∼20 seconds), this suggests a substantial portion of chromis foraging behaviour is constrained by occupying positions in the water column with reduced opportunities for feeding. This pattern is consistent with the concept of a “landscape of fear” [19,24], where prey modify habitat use based on spatial variation in perceived predation risk. While reactive non-consumptive effects result in strong but brief reductions in feeding, proactive responses may exert a more continuous and cumulative influence on energy intake. Chromis may forgo higher foraging rewards higher in the water column in favour of safer, but less productive zones, incurring a steady cost that may outweigh the short-term effects of predator disturbances. These findings underscore the importance of both immediate and anticipatory risk responses in shaping energy transfer from pelagic sources to reef-associated consumers.

### Future applications

Our pipeline opens the door to a wide range of ecological applications where direct measurement of interaction rates has previously been infeasible. Within coral reef ecosystems, it could be used to quantify how rates of planktivory vary across reef zones (e.g., fore reef vs. back reef), between different coral morphologies, or over diel cycles. Such measurements could help test hypotheses about how physical habitat structure modulates foraging behaviour and energy flow. When paired with reef-wide surveys of fish abundance, targeted behavioural measurements could also be used to scale up from individual-level feeding to community-level estimates of total planktivory across a reefscape. In addition, by measuring feeding rates across a range of local fish densities, the tool could be used to explore density dependence in planktivory under natural conditions—a question that has been difficult to address in situ. The ram-jaw feeding behaviour of *Chromis multilineata* is widespread among planktivorous damselfishes [53], suggesting that this approach could be readily extended to numerous other species within the Pomacentridae family.

While we implemented the approach using a specific set of object detection, tracking, and classification models, the pipeline itself is modular and highly adaptable. Different model architectures or training strategies can be substituted at each stage to tailor the system to new species, behaviours, or environments. The pipeline can be applied with minimal modification to quantify any interaction where the relevant behaviours are visually recognisable in static frames, such as foraging strikes, courtship displays, or aggressive postures. In systems where the interacting species are both visible and sufficiently large, the pipeline could be extended to classify interactions based on the proximity and identity of two individuals, in addition to behavioural cues. For interactions that are not visually discernible from single frames and require temporal context, video-based deep learning approaches, such as anchor-free temporal action detection models or transformer-based models, may be required to detect interactions [54]. Nonetheless, the core pipeline we present, combining tracking, frame-based classification, and bias correction, offers a robust, generalizable framework for quantifying a diverse range of visually identifiable ecological interactions.

### Limitations

An important limitation of our approach in the context of quantifying planktivory is that, due to the large size disparity between planktivorous fish and their zooplankton prey, our pipeline captures only the behaviour of the predator, not the identity or size of the prey being consumed. While previous stomach content analyses suggest that brown chromis feed primarily on copepods (comprising ∼87% of their diet), copepod species can vary significantly in body size and energy content. This variation makes it difficult to directly convert feeding event rates into biomass or energy flux without additional information on prey identity. To translate behavioural data into estimates of energy transfer or trophic flux, our approach would need to be complemented by concurrent measurements of prey availability and composition, such as paired plankton tows, gut content analysis, or emerging optical tools capable of resolving prey in situ [55,56]. Incorporating such data would allow for converting observed feeding rates to biomass or energy fluxes, improving estimates of the ecological consequences of planktivory across reef environments. Importantly, this limitation is specific to predator–prey interactions involving highly size-mismatched species. In other systems, such as mutualisms, competition, or aggression, where interacting species are of comparable size, the identities of both participants may be directly observable, and this constraint may not apply.

Because our classifier detects prey capture attempts rather than successful ingestion, feeding rates may be overestimated if capture success is low. However, for ram-feeding species like *Chromis multilineata*, prey capture success is generally high (∼90%) even against evasive copepods [33], and is therefore unlikely to introduce substantial bias into feeding rate estimates or comparisons across time or environmental gradients.

### Conclusions

Our study demonstrates how machine-learning-based computer vision can be used to directly quantify the rate of ecological interactions in the field, overcoming long-standing challenges in measuring dynamic processes like planktivory. By combining automated detection, tracking, behavioural classification and bias correction, our approach enables scalable, high-resolution estimates of feeding rates in natural conditions. As automated computer vision tools continue to advance, they offer powerful new opportunities to link behaviour with ecological function, resolving how individual behaviours scale up to shape ecosystem dynamics.

## Acknowledgements

We thank Lena Faber and Luna Buwalda for their assistance with field work, and Andrew Hein and Abigail Grassick for their feedback on the machine-learning pipeline.

## Data availability statement

The data that support the findings of this study are openly available in “figshare” at https://figshare.com/articles/dataset/Automated_interaction_rates/29069525, reference number 29069525.

## Authorship Statement

BM collected the data. MS and BM developed the pipeline and conducted the analysis. MS wrote the first draft of the manuscript and both authors contributed to further revisions.

