## Supplementary for "Automated quantification of ecological interactions from video"

#### Supplementary Figure 1

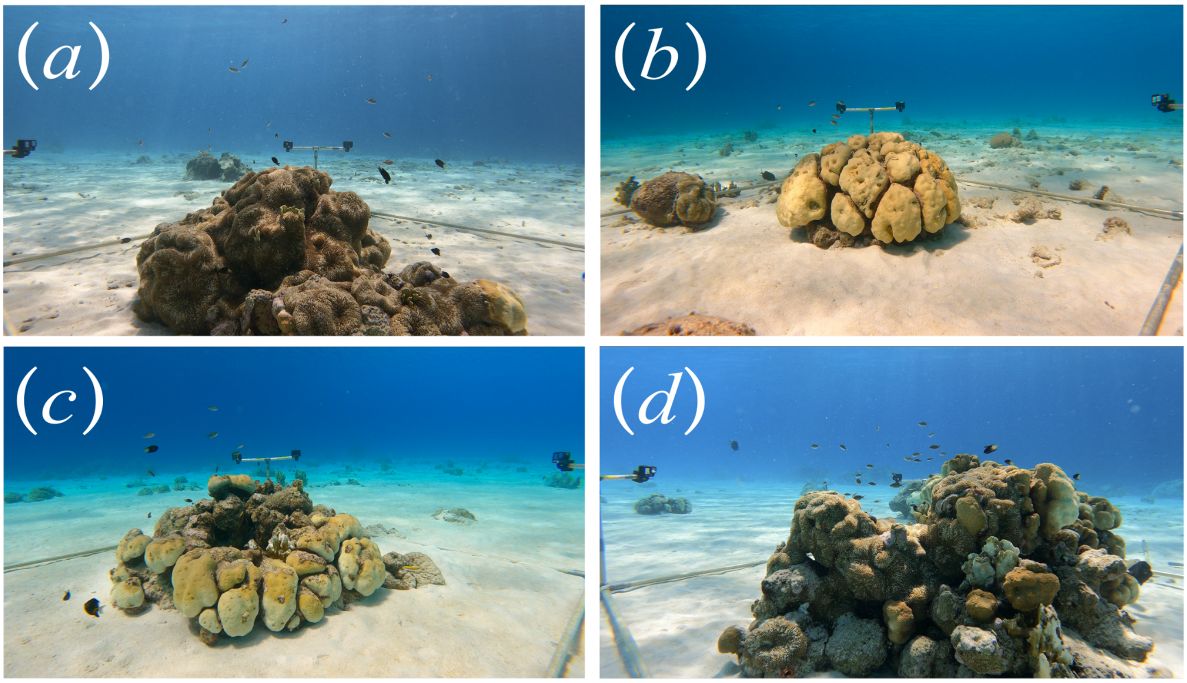


**Supplementary Figure 1. Coral-head study sites.** (a) Study site 1, (b) study site 2, (c) study site 3, and (d) study site 4 can be seen from the perspective of one camera in the underwater video-recording set-up.

#### Supplementary Figure 2

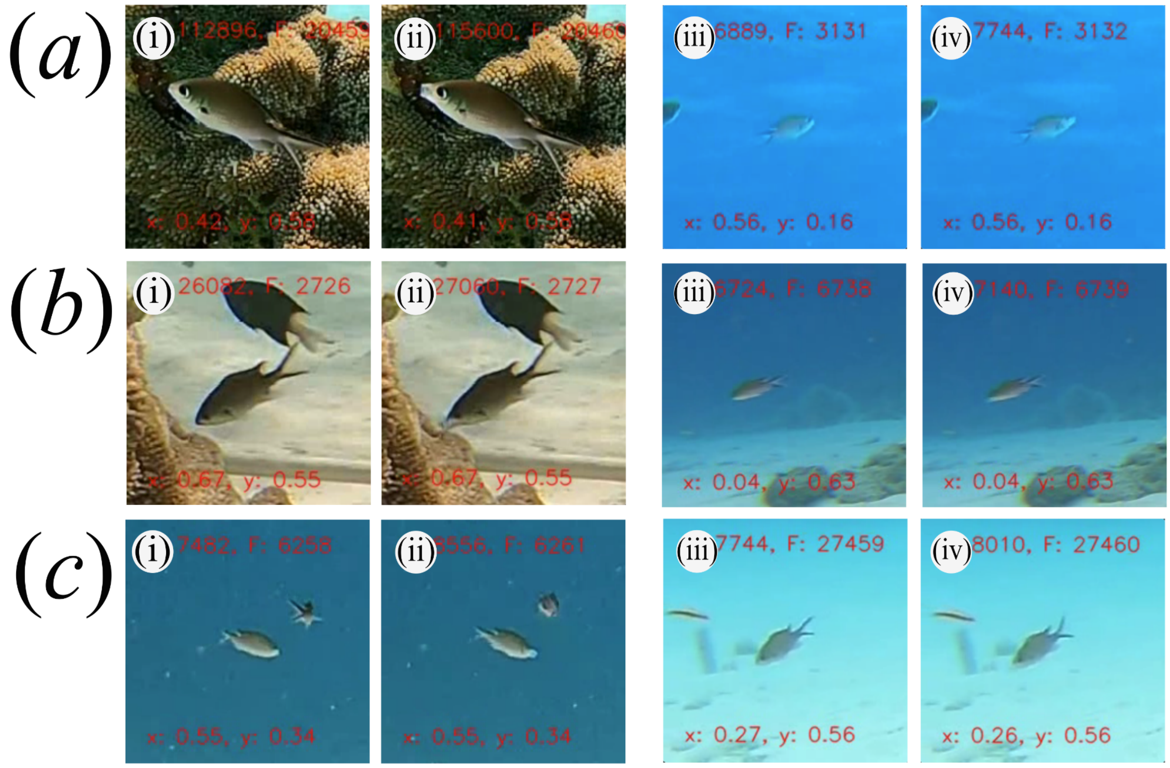


**Supplementary Figure 2. Feeding behaviour of brown chromis across different frame-associated factor conditions.** (a) (i-ii) large and (iii-iv) small pixel size of the tracked object or "bounding box area"; (b) (i-ii) centre and (iii-iv) edge positions of the tracked object with respect to the camera view or "coordinates"; (c) (i-ii) high and (iii-iv) low contrast between feeding behaviour and background or "red, green and blue background colour contents". For each factor (a, b & c), tracked chromis with high feeding behaviour visibility (i-ii) and low feeding behaviour visibility (iii-iv) are shown. For both high and low feeding behaviour visibility in each frame-associated factor, the second image (ii & iv) shows the brown chromis fish feeding, and the first image shows the frame just before that (i & iii).

### Supplementary Table 1

**Supplementary Table 1. Logistic regression models for the estimation of image classification model recall and precision based on frame-associated factors.** Logistic regression models used to calculate precision and recall for the image classification model on "Feeding" and "Not Feeding Mouth Visible" classes, based on frame-associated factors, are presented. Frame-associated factors include: average bounding box area (bb_area); background red (r), green (g), and blue colour content (b); bounding box "y" coordinates (y); and bounding box "x" coordinates, transformed to represent the number of pixels from the centre x coordinate (transformed_x). These equations were used to calculate the expected recall and precision for each class over a given period by inputting the relevant mean values of the frame-associated factors across all frames contributing to the feeding rate estimate. Details on how the models were obtained can be found in the Methods section "Model evaluation and bias correction".

| Class | Performance Metric Estimated | Logistic Regression Model |
|---|---|---|
| Feeding | Precision | logit(P) = (-1.0543) + (0.3857*log(bb_area)) + (-0.0078*r) |
| Feeding | Recall | logit(P) = (-7.9686) + (1.2174*log(bb_area)) + (-0.0265*g) + (0.0179*b) + (-0.0005*transformed_x) |
| Not Feeding Mouth Visible | Precision | logit(P) = (-0.6602) + (0.5943*log(bb_area)) + (-0.0017*r) + (-0.0052*g) + (0.0039*b) + (0.0001*y) + (-0.0005*transformed_x) |
| Not Feeding Mouth Visible | Recall | logit(P) = (-12.1735) + (2.0282*log(bb_area)) + (-0.0031*r) + (-0.0033*b) + (-0.0003*y) + (-0.0007*transformed_x) |

#### Supplementary Table 2

**Supplementary Table 2. Image classification model performance metrics ("test" dataset).** Precision ("Precision", equal to true positives/total predicted positives) and recall (true positives/total positives present in the dataset) for each of the individual brown chromis behaviour classes are presented for the image classification model. These metrics were obtained by evaluation on the "test" dataset, comprising 24 837 images, of which 557, 5 507, 584 and 18 189 belonged to the "feeding", "mouth not visible", "non-chromis" and "not feeding mouth visible" classes, respectively.

| Individual chromis behaviour | Precision | Recall |
|---|---|---|
| Feeding | 0.935 | 0.698 |
| Mouth not visible | 0.870 | 0.866 |
| Not feeding mouth visible | 0.989 | 0.938 |
| Non-chromis | 0.950 | 0.961 |

#### Supplementary Table 3

**Supplementary Table 3. Image classification model performance metrics on brown chromis tracks without exclusion of small bounding box areas.** Model performance metrics (as described in Table 2) were obtained by comparison of model predictions and manual annotations for 1 998 randomly selected and individual brown chromis tracks, without exclusion of tracks with bounding box sizes lower than the 1 451 pixel threshold.

| Individual chromis behaviour | Precision | Recall |
|---|---|---|
| Feeding | 0.858 | 0.605 |
| Mouth not visible | 0.493 | 0.872 |
| Not feeding mouth visible | 0.969 | 0.815 |
| Non-chromis | 0.519 | 0.741 |

#### Supplementary Table 4

**Supplementary Table 4. Image classification model performance metrics on bounding box area-filtered individual brown chromis tracks.** Model performance metrics (as described in Table 2) were obtained by comparison of model predictions and manual annotations for 1 998 randomly selected and individual brown chromis tracks, filtered to contain only frames with bounding box areas above the 1 451 pixel threshold.

| Individual chromis behaviour | Precision | Recall |
|---|---|---|
| Feeding | 0.865 | 0.761 |
| Mouth not visible | 0.506 | 0.795 |
| Not feeding mouth visible | 0.979 | 0.932 |
| Non-chromis | 0.941 | 0.910 |
